# Mutation in position of 32 (G>U) of S2M differentiate human SARS-CoV2 from Bat Coronavirus

**DOI:** 10.1101/2020.09.02.280529

**Authors:** Majid Vahed, Mohammad Vahed, Aaron Sweeney, Farshad H Shirazi, Mehdi Mirsaeidi

## Abstract

The new Severe Acute Respiratory Syndrome Coronavirus 2 (SARS-CoV-2) is a zoonotic pathogen that has rapidly mutated and become transmissible to humans. There is little existing data on the mutations in SARS-CoV-2 and the impact of these polymorphisms on its transmission and viral load. In this study, the SARS-CoV-2 genomic sequence was analyzed to identify variants within the 3’UTR region of its cis-regulatory RNA elements. A 43-nucleotide genetic element with a highly conserved stem-loop II-like motif (S2M), was discovered. The research revealed 32 G>U and 16 G>U/A mutations located within the S2M sequence in human SARS-CoV-2 models. These polymorphisms appear to make the S2M secondary and tertiary structures in human SARS-CoV-2 models less stable when compared to the S2M structures of bat/pangolin models. This grants the RNA structures more flexibility, which could be one of its escape mechanisms from host defenses or facilitate its entry into host proteins and enzymes. While this S2M sequence may not be omnipresent across all human SARS-CoV-2 models, when present, its sequence is always highly conserved. It may be used as a potential target for the development of vaccines and therapeutic agents.

## Introduction

The emergence of new viral pathogens is a danger to public health (1). Three emerging pathogenic coronaviruses (CoVs), Severe Acute Respiratory Syndrome Coronavirus (SARS-CoV-1), Middle East Respiratory Syndrome Coronavirus (MERS-CoV), and a newly identified CoV (SARS-CoV-2), are zoonotic viruses, which utilize bats as their natural reservoir. They are then transmitted through intermediate hosts, eventually infecting humans. The pressure on host selection of SARS-CoV-2 in human models will have an effect on long term conservation of mutations that enhance its transmissibility (2). However, the determinants regulating strong trans-species evolution remain unknown due to challenges in recognizing viral precursors and animal reservoirs (3). A 43-nucleotide genetic element with a highly conserved stem-loop II-like motif (S2M) has been reported in four groups of positive single stranded RNA viruses; Astroviridae, Picornaviridae, Caliciviridae, and Coronaviridae (4).

The significance of S2M sequences in viral strains remains to be determined. However, it appears to be important to viral transmissibility and will likely be found in future emergent coronaviruses (5). Given that S2M is a highly conserved component of coronaviruses viral genome and is absent from the human genome, it could become a potential target for antiviral drug therapy (6). This study was performed in order to analyze coronavirus genomic sequences isolated from human, bat, and pangolin models in order to identify variants within the structured cis-regulatory RNA elements in the 3’UTR region, including the S2M loop.

## 2. Methods

### 2.1. Calculation of Allele Frequency and Occupancy

The SARS-CoV-2 isolated from patients on a cruise ship in Japan (NCBI GenBank Accession Number LC528232.1) was used as a reference genome. Identifying functional RNA motifs and elements was performed using RegRNA 2.0 tools (filtered to human settings) (7).

The S2M motifs (43 nucleotides long) were aligned for bat/pangolin models (Fig. 1 (a)) and for human models SARS-COVs (Fig. 1(b)). A G to U amino acid transfer at position 32 and a G to U/A amino acid transfer as position 16 (Fig. 1 (b)) were found in human SARS-CoV-2. This profile was used to search for all viral sequences in GenBank, while using different combinations of 32 G>U and 16 G>U/A nucleotide substitutions that occur in a conserved region within 3’ UTR, known as the S2M motif.

**Figure 1.**
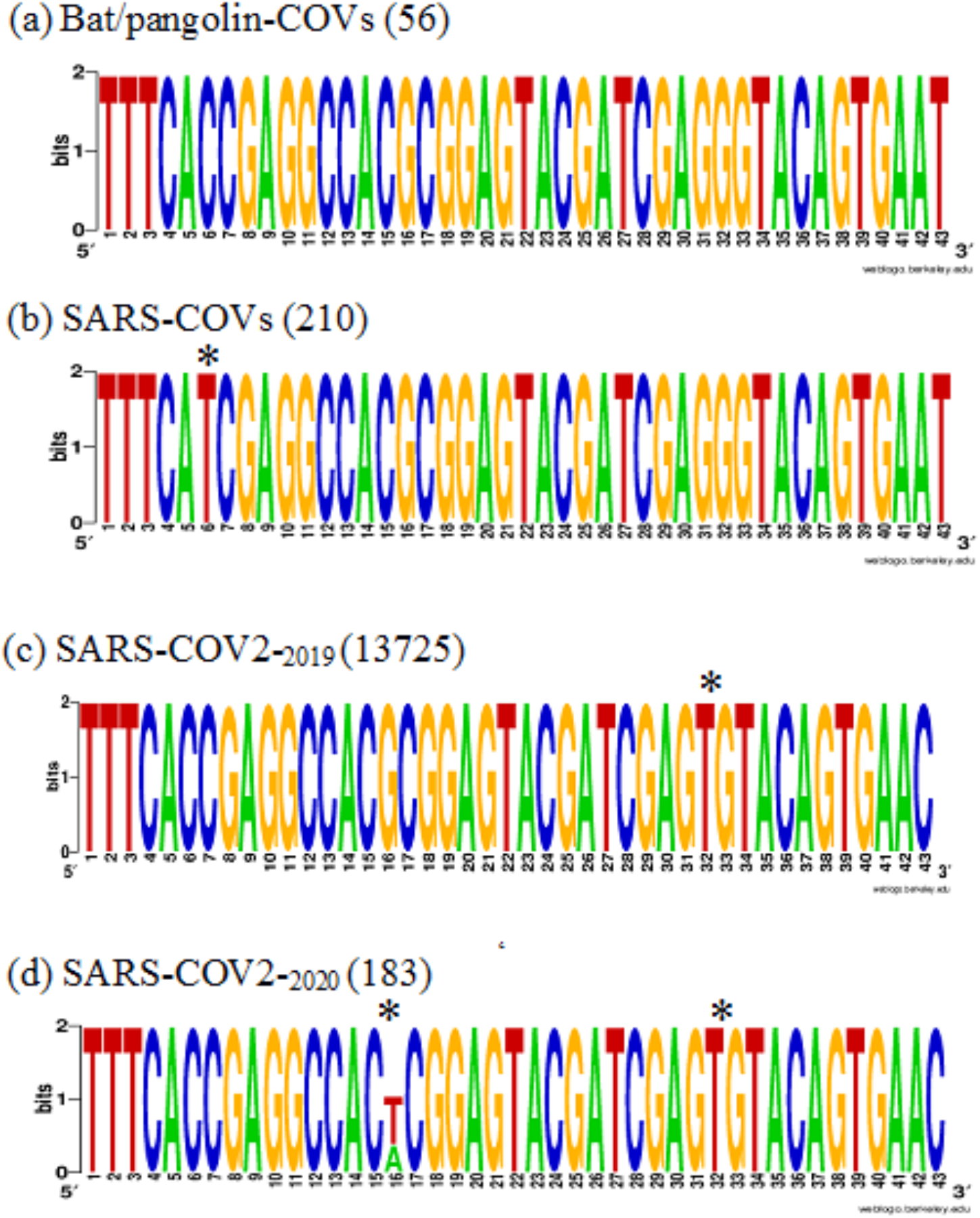
Multiple sequence alignment logos of selected S2M motifs. (a) BLASTn search of the NCBI database for G32 S2M motifs (b) Human U32 S2M coronavirus motifs. Nucleotide letter heights indicate their frequency at the G32 position in bat and pangolin S2M hairpins.

### 2.2. Quantifying Similarities between Motifs and Structures

RNA sequences were aligned using 3’UTR (29,543–29,903), with queries in a BLASTn search of the NCBI database for S2M motifs(8, 9). The 3’ UTR stem-loop structures were determined by using the RNAfold web server (http://rna.tbi.univie.ac.at/forna/). Additionally, sequence and structural information of astrovirus S2M motifs was integrated for RNAfold design to analyze a potential zoonotic precursor for coronaviruses from a hypothetical infectious sequence (10). The PDB structure for S2M was downloaded from the Protein Data Bank (http://www.rcsb.org/pdb) (PDB ID: 1XJR) (6). All of the structures were visualized by PyMOL (11) and analysis was performed as presented in previous studies (12, 13). RNAComposer was used to model the three dimensional (3D) RNA structures (14, 15). Clustal Omega was used to apply mBed algorithms for guide trees. ClustalW alignment tools were used to execute multiple sequence alignments (16). The S2M RNA sequences were compared between members of the coronavirus family (bat and pangolin CoVs, SARS-CoV-1, and SARS-CoV-2) as well as the astrovirus family (avian, porcine, bovine, chicken, turkey, ovine, mink, tiger, human and mamastrovirus). The S2M motifs were screened with miRBase tools in order to identify potential miRNA binding sites (17).

### 2.3. Identification of Single Nucleotide Substitutions in the SARS-CoV-2 Genome

Only alignment positions which harbored defined A/T/G/C residues in 95% of their genomes were considered for nucleotide substitutions (18). In order to be graphically represented, sequence logos of the selected S2M motifs were then constructed using WEBLOGO (19–21).

A nucleotide BLAST search of these sequence motifs was performed on the NCBI portal. S2M containing sequences were identified for the following sequence: 5’-TTTCA (G>U/A)CGAGGCCACGCGGAGTACGATCGAG(G>U)GTACAGTGAA(U>C)-3’.

This text string profile was used to search for all the viral sequences in GenBank available as of August 14^th^, 2020, while using different combinations of the aforementioned nucleotide substitutions. A phylogenetic tree was created by generating a genomic epidemiological map of SARS-CoV-2 at position 29724 (S2M) using NextStrain tools (https://nextstrain.org/) (22). The sequences of the different isolates were obtained from GISAID (https://www.gisaid.org/) (23).

## 3. Results

### 3.1. Calculation of Allele Frequency and Occupancy

Position 32 of S2M is a critical point of the sequence that is specific to the human coronaviruses and variable for the bat/pangolin coronaviruses (Fig. 1 (a,c)). A second polymorphism appears in human SARS-CoV-2 models at position 16 G>U/A (Fig. 1 (d)). A previous polymorphism was found in SARS-CoV-1 at position 6 C>U (Fig. 1 (b)).

### 3.2. Calculation of Cis-Regulatory RNA structures and Minimum Free Energy

Rapidly emerging variations within the cis-regulatory RNA structures of the SARS-COV-2 genome were analyzed and characterized. The hotspot of S2M motif mutations was found to occur at position 32 (G>U), where a single mutation occurred between bat/pangolin and human stem-loop structures.

The impact of 32 G>U mutations decreased RNA secondary structure stability in both MFE (−8.80 kcal/mol vs. −6.20 kcal/mol) and centroid structures (−8.80 kcal/mol vs. −0.80 kcal/mol) when using RNAfold calculations (Fig. 2). The 16 G>U mutations in SARS-CoV-2 improved RNA secondary structure stability in both minimum free energy (MFE) (−8.80 kcal/mol vs. −11.70 kcal/mol) and centroid structures (−8.80 kcal/mol vs. −9.90 kcal/mol) when using RNAfold calculations (Fig. 2).

**Figure 2.**
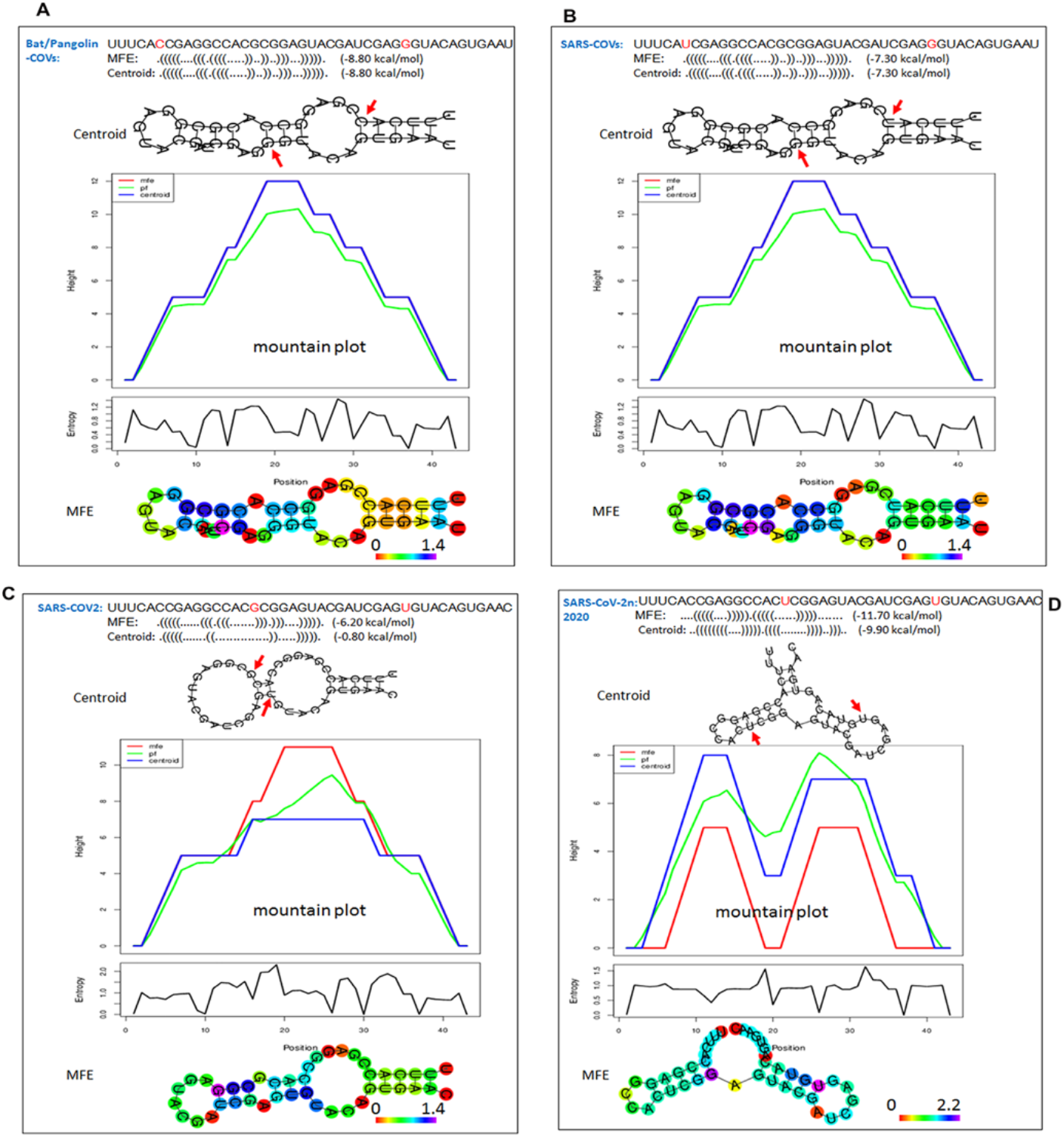
The minimum free energy structure (MFE) and centroid structures for S2M motifs. Base pairs are colored according to base pair probabilities computed with thermodynamic parameters using the Vienna Program [12,13]. The mountain plot is an xy-graph that represents secondary structures including MFE, thermodynamic ensemble of RNA (pf), and centroid structures in a plot of height versus position. “MFE” represents the minimum free energy structure, “pf” indicates partition function and “centroid” represents the centroid structure. The height m (k) is given by the number of base pairs enclosed at position k. Three curves are shown: the MFE structure (red), the pairing probabilities (black) and a positional entropy curve (green). Well-defined regions are identified by low entropy. (b) Structural changes where G is substituted by U. The positional entropy is coded by hues ranging from red (low entropy, well-defined) to violet (high entropy, ill-defined). The lower panel depicts entropy vs position.

Interestingly, the MFE and centroid secondary S2M structures of SARS-CoV-2__2020_ are significantly different from human and bat/pangolin coronaviruses. As shown in Fig. 2, the G>U base changes at residues 16 and 32 influence the folding of hairpin loops.

Additional analysis of canonical and MFE S2M steam-loop structures is shown by the dot plot (Fig. 3). By superimposing various mountain plots, the bat/pangolin S2M secondary structures appear to show equal distribution on both sides. However, the SARS-CoV-2 S2M secondary structures shows non-uniform distribution (Fig. 3). The closer the two curves, the better is the defined diffraction of the S2M structures in human and bat/pangolin coronaviruses.

**Figure 3.**
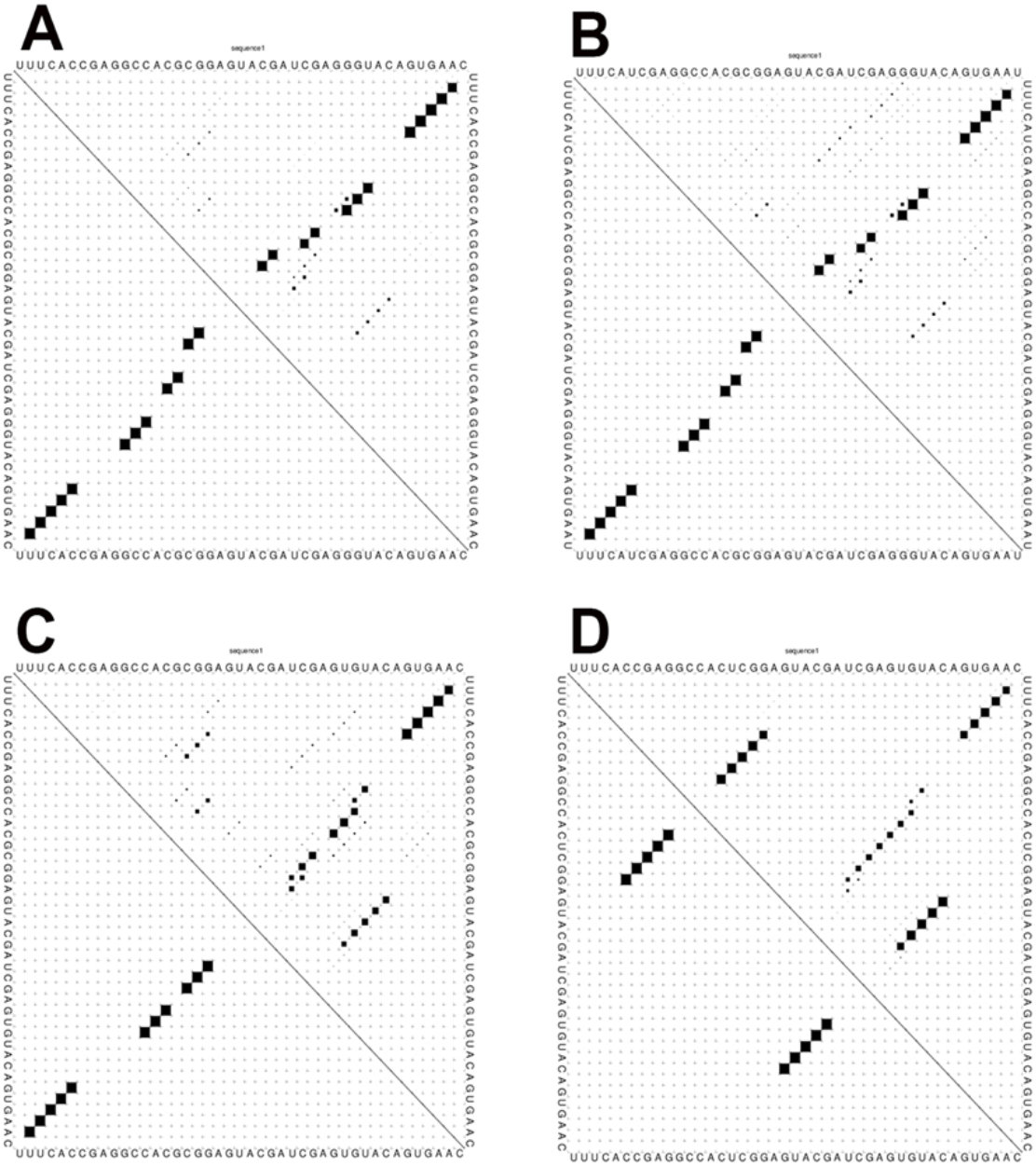
Dot plot produced by RNAfold. The MFE structure is represented in the lower left triangle. The probability of all possible base pairs is represented in the top right triangle. The area of each dot is proportional to the pairing probability of the base pairs. (a) Human coronavirus S2M, (b) Bat/pangolin coronavirus S2M. The resulting plot depicts three curves. The red curve shows two peaks derived from the MFE structure, the black curve demonstrates pairing probabilities, and the green curve indicates positional entropy. Every dot symbolizes a base pair. The size of the dots in the upper right triangle is proportional to the respective base pairing probability. Changes in the MFE structure of human coronavirus S2M in the lower left triangle.

The centroid structures of RNA S2M sequences with minimal base-pair distance are significantly different in human MFE structures of astroviruses, and are similar in other astroviruses (Fig. S3, S4).

### 3.3. The Tertiary Structures of SARS S2M RNA

3D structures of S2M sequences of coronaviruses in human and bat/pangolin models were created based on the secondary structure in the dot-bracket format (Fig. 4). The mutations at position 16 and 32 (G>U) impact tertiary structures and consequently cause conformational changes. The human stem-loop structure is bent to the right side when compared to the stem-loop structure in the bat/pangolin coronavirus (Fig. 5). The conformational changes within the S2M loop may affect its binding to host proteins and enzymes.

**Figure 4.**
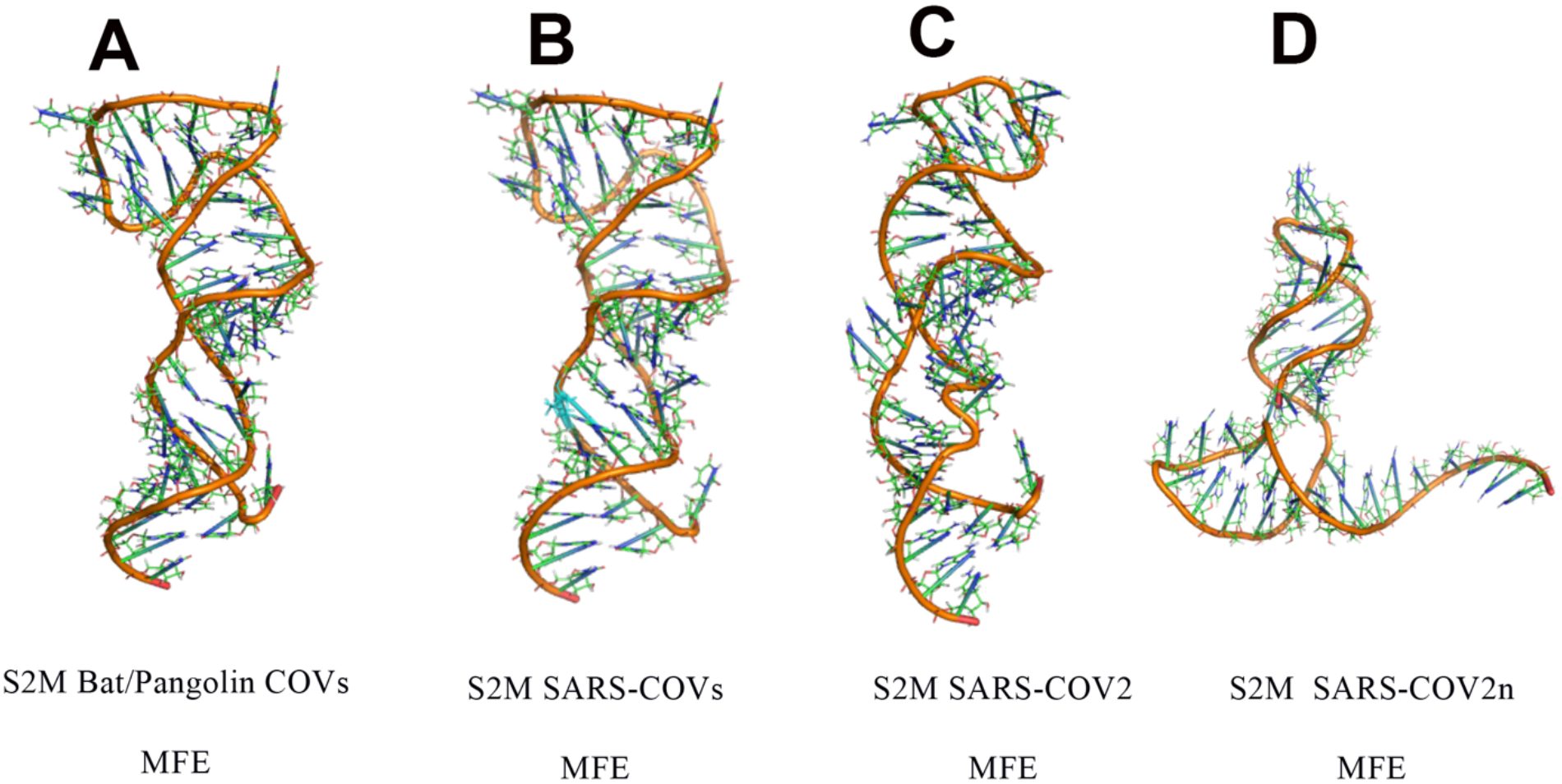
Tertiary structures of SARS-CoV-2 S2M RNA. Phylogenetic comparisons of S2M sequences from (a) Bat/Pangolin coronaviruses, (b) Human coronavirus MFE structure and (c,d) Human coronavirus canonical structure.

**Figure 5.**
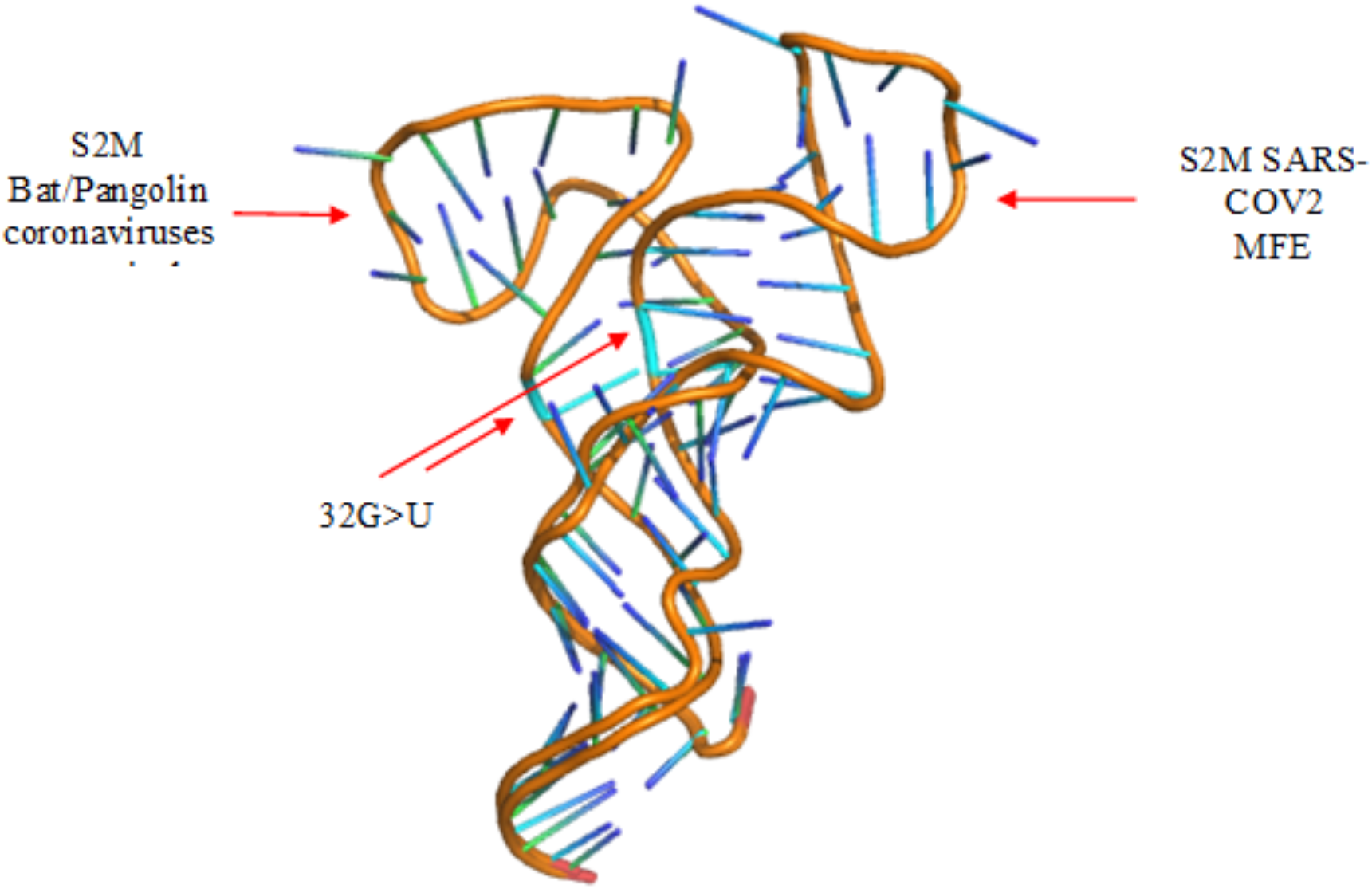
Superposition of tertiary structures of SARS-CoV-2 S2M RNA from bat and pangolin coronaviruses and human coronavirus MFE structure.

### 3.4. Alignment and Clustering of S2M Stem-Forming Elements and Columns

ClustalW multiple sequence alignment was used to align secondary structures. The great majority of the sequences were able to fold into the canonical S2M stem-loop structure. Clusters of S2M sequences are highlighted. The clustering trees of coronavirus (Fig. 6) and astrovirus (Fig. S2) are displayed. The 32 G>U mutation introduces changes in Watson–Crick pairing, with resultant changes in the predicted secondary structures. ClustalW multiple sequence alignment reveals that 32 G>U base-pairing exhibits complementary changes to the secondary structures.

**Figure 6:**
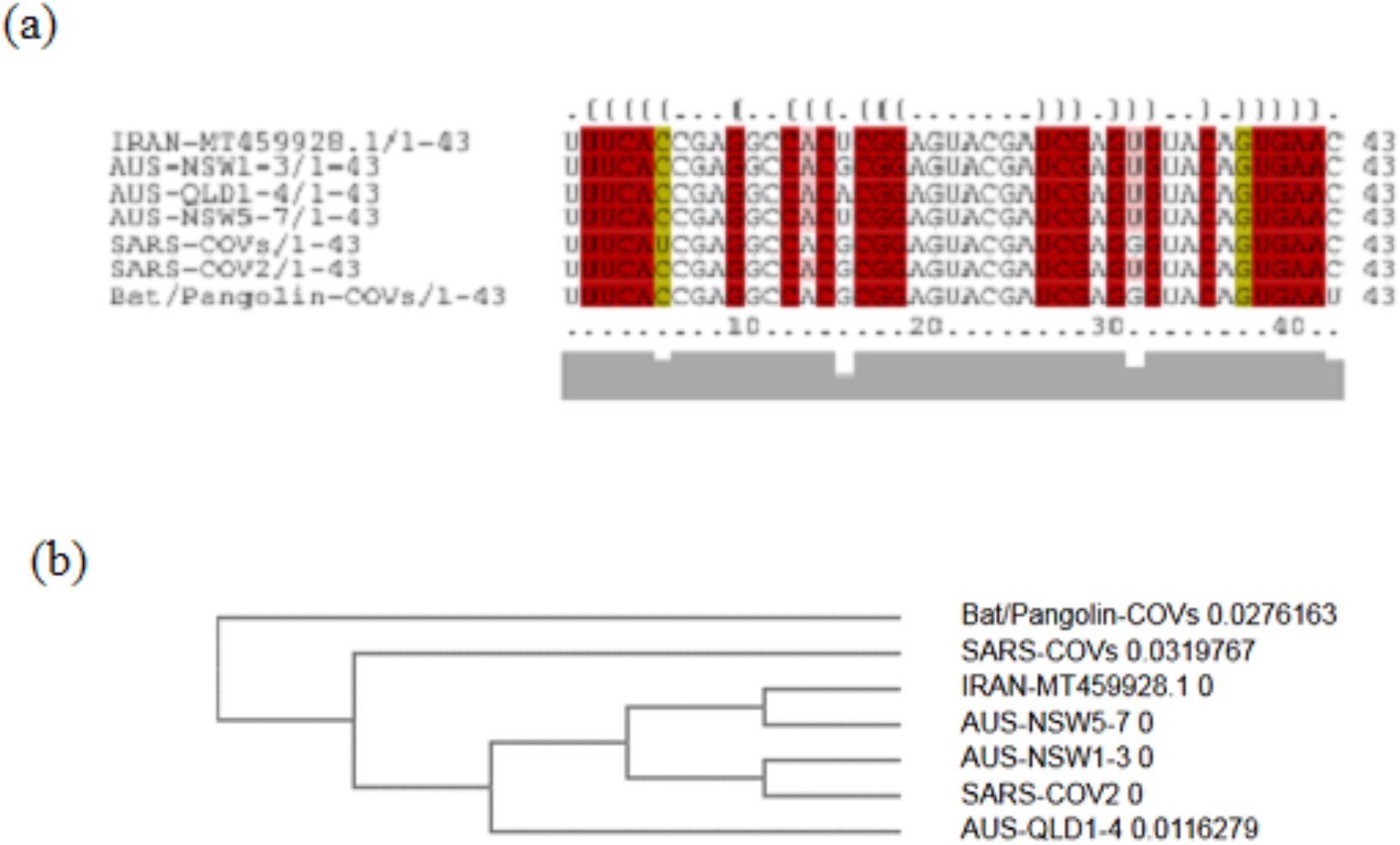
Coronavirus S2M sequence motifs. (a) Alignment of stem-forming elements and columns for each genotype for sequence representation and stem-loop structure representation. Lines above the alignment indicate stem-forming elements. Columns with Watson–Crick pairing changes for nucleotide positions have been color-coded. (b) ClustalW multiple sequence alignment trees display of coronavirus.

### 3.5. Potential Interaction of S2M motifs and Human miRNAs

Mirbase was used to screen human miRNA that could target S2M sequences. Additional focus was put on miRNAs that have been reported as components of anti-viral miRNA-mediated defense [28]. This study identified two potential binding sites within the S2M sequences of bat/pangolin and SARS-CoV-1: hsa-miR-1304-3p & hsa-miR-1307-3p. Only one potential binding site was found within the S2M of Australian and Iranian SARS-CoV-2 samples: hsa-miR-1307-3p (Fig. 7).

**Figure 7:**
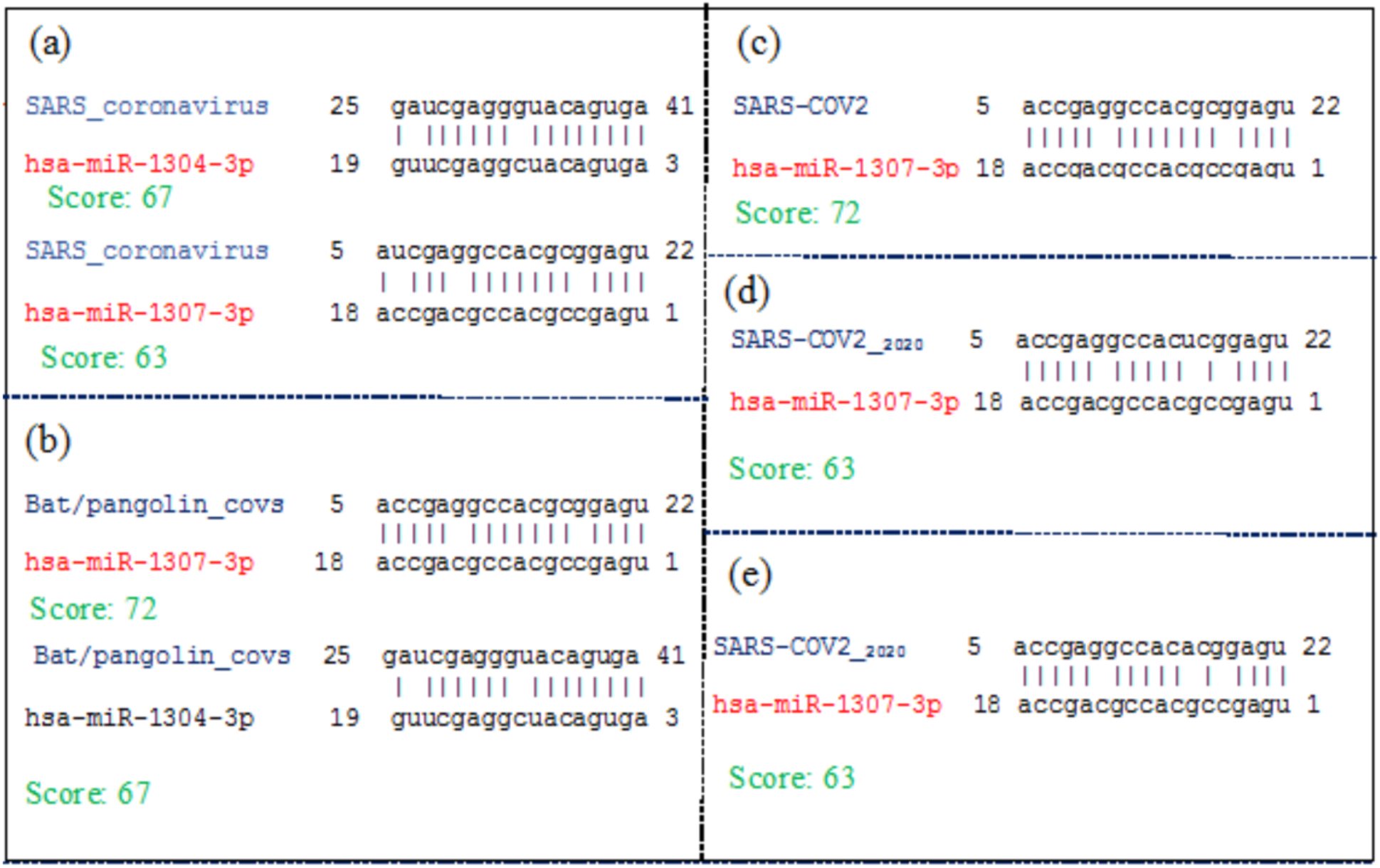
The predicted human miRNA binding sites within different regions of S2M sequences. (a) miRNA binding sites within the S2M of SARS-CoV-1 (b) Bat/pangolin coronavirus. (c, d, e) The miRNA binding sites within the S2M of SARS-CoV-2.

### 3.6. Earlier mutations of 16 G>U/A S2M motifs

While analyzing all available sequences available up to August 14^th^ 2020, a 16 G>U/A S2M motif mutation was identified at position 29742 of SARS-CoV-2_-2020_ in Iran, Australia, Taiwan, Sir Lanka, Bahrain, USA, Georgia, Bangladesh, Norway, Hong-Kong, Germany and Turkey (Fig. 8 and 9).

**Figure 8:**
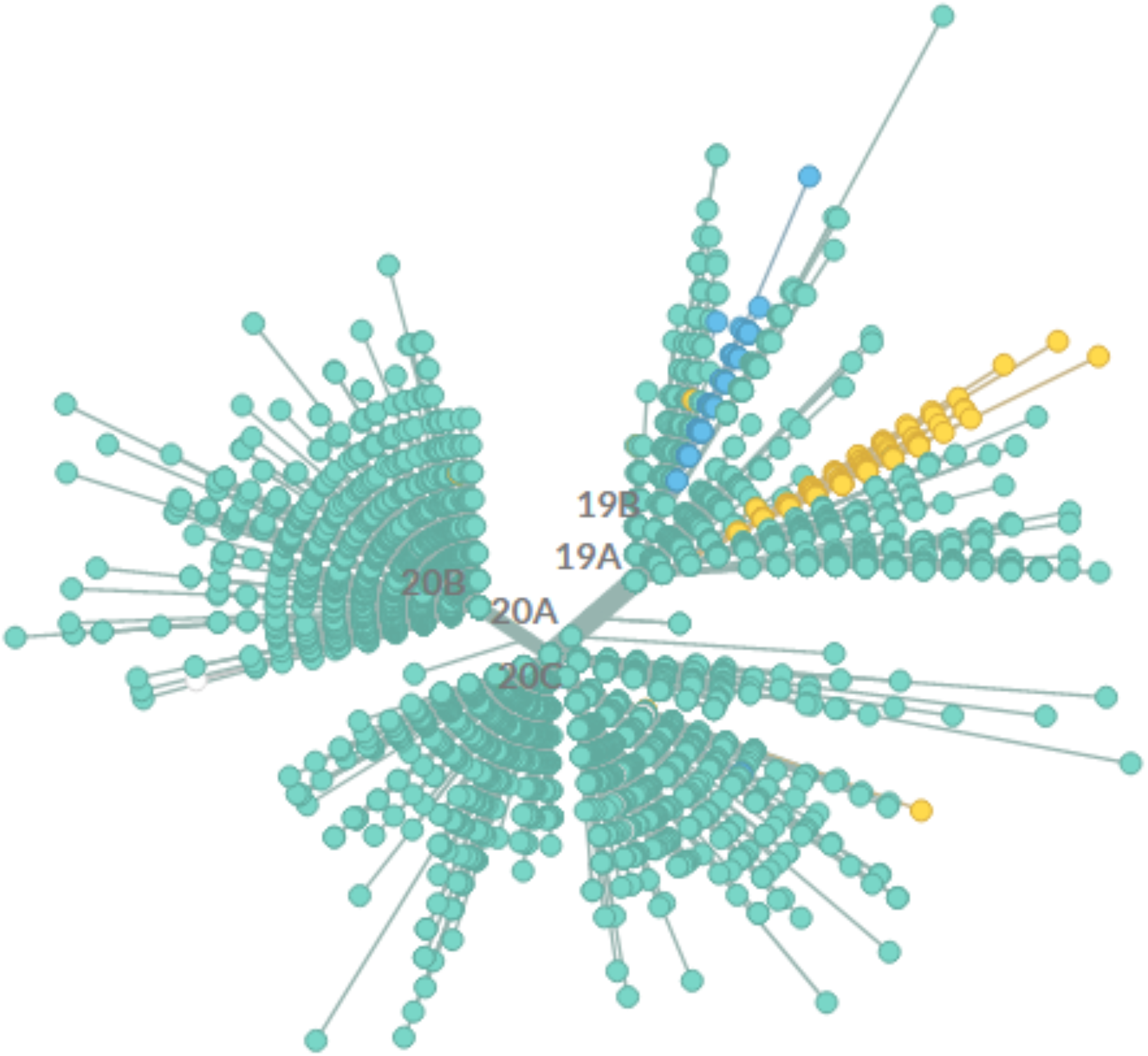
The position of conserved mutations in a phylogenetic graph obtained from Nextstraindatabase. Picture captured on August 14st, 05: 17 AM.

**Figure 9:**
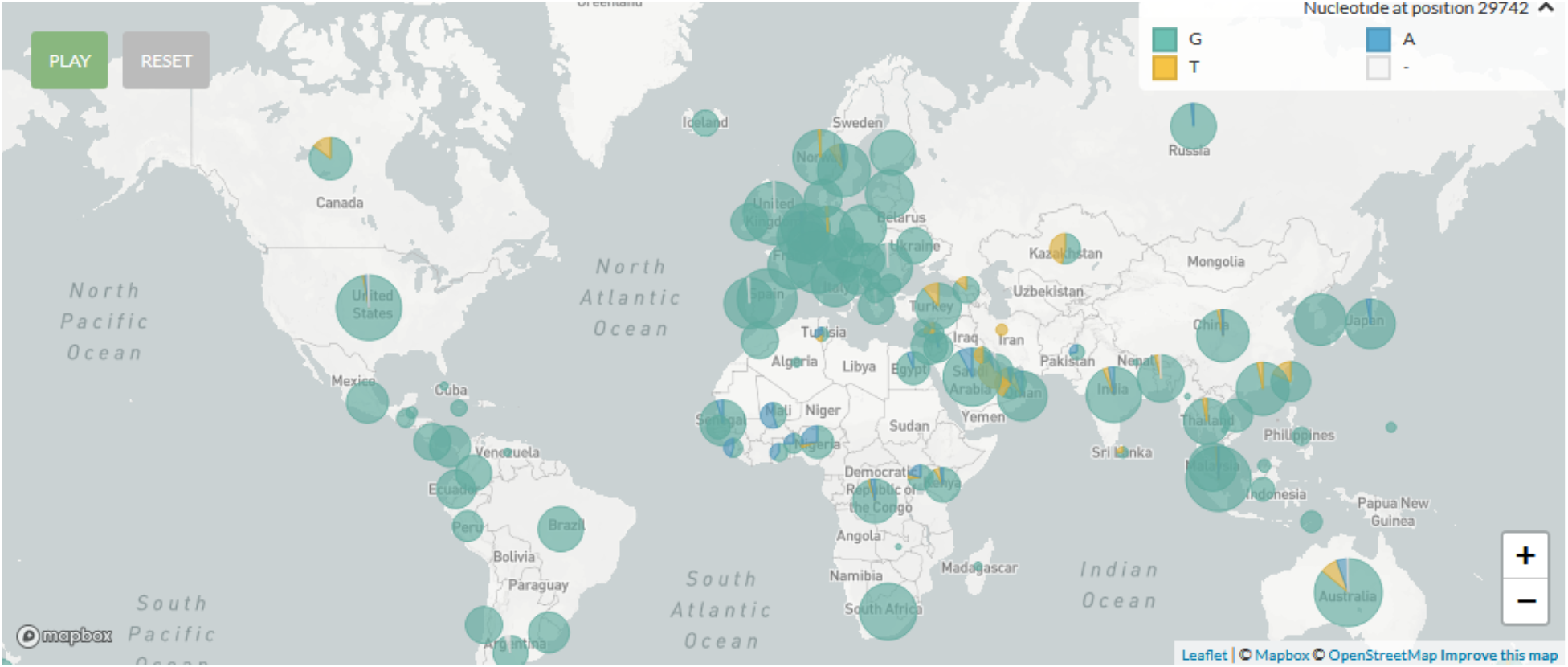
Geographical map of SARS-CoV-2 at position 29742 (S2M (16)).

## 4. Discussion

The current study found that a consistent G>U mutation at the 32 position, of the 43 nucleotide, long S2M sequence of SARS-CoV-2. This mutation has not been found in any bat or pangolin CoV strains. We concluded that transmissibility to human from bat/pangolin was related to this mutation.

The MFE^1^ and centroid secondary structures were found to be significantly different in SARS-CoV-2 sequences in comparison to bat/pangolin coronavirus sequences. This appears to be due to the mutation occurring at position 32 of S2M. There were also significant differences in the MFE^1^ and centroid structures between the human SARS-COV-2 and SARS-COV-2_2020 S2M sequences. The MFE and centroid secondary structures were observed to be similar in S2M bat/pangolin coronavirus and SARS-COVs.

Ancestral recombination and genetic similarity between coronaviruses infecting bats/pangolins and humans may have played a key role in the evolution of the strain that led to the introduction of SARS-CoV-2 in humans (24).

The analysis of Australian SARS-CoV-2 sequences revealed clustering of mutations in S2M secondary structures across coronaviruses and astroviruses. These new changes within the S2M sequences and structures could be the virus’ escape mechanism from host defense systems. The SARS S2M secondary structures containing G (16), and G (32) are conserved in Bat/pangolin related coronaviruses. A/U (16) and U (32) mutations are found in SARS-COV-2-2020. These changes create a less stable RNA structure in SARS-CoV-2, thereby granting it more flexibility, which could be one of its escape mechanisms from host defenses or facilitate its entry into host proteins and enzymes

Based on the structural similarities between micro RNA hairpins implicated in gene regulation, it is possible that S2M functions through an RNA interference-like mechanism, and possibly targeting homologous sequence loci in the infected organisms (25).

32G>U mutations may change the RNA secondary structures that are critical to viral reproduction or interfere with sequences targeted by host miRNAs. While screening for human miRNA targets on S2M motifs, two targets sequences were found within the bat/pangolin and SARS-CoV-1 S2M and one within the SARS-CoV-2. The presence of host miRNA targets within S2M motifs may be crucial for host selection and anti-viral miRNA-mediated defense. The S2M of SARS-CoV-2 may promote its viability and infectivity.

It is likely that the S2M sequences of (+) ssRNA viruses are still active and will continue to affect these viruses’ evolution [5]. The 16 G>U/A and 32 G>U nucleotide changes in the S2M sequence of SARS-CoV-2 render it a target for one human miRNA, hsa-miR-1307-3p. However, bat/pangolin coronavirus and SARS-CoV-1 S2M sequences are targetable by two human miRNAs hsa-miR-1307-3p and hsa-miR-1304-3p. As such, only one human miRNA is capable of targeting the S2M sequence of SARS-CoV-2 and affecting its viral replication.

## 5. Conclusions

The study examined S2M mutations able to impact the cis-regulatory elements in the SARS-CoV-2 genome. While the evolutionary and functional origin of S2M has yet to be discovered, its presence across even distantly related viruses insinuates that the sequence is important for viral transmission. These findings provide insight into the significance of viral RNA structures and introduce S2M as a potential target for development of vaccines and therapeutic agents.

## Acknowledgments

Authors would like to that Nextstrain for providing a real-time snapshot of evolving SARS-CoV2.

