## Supplementary material for "Mutation in position of 32 (G>U) of S2M differentiate human SARS-CoV2 from Bat Coronavirus": NA

**Affiliations:**
[1]Pharmaceutical Sciences Research Center, Shahid Beheshti University of Medical Sciences, Niayesh Highway, Valiasr Ave, Tehran, Iran

[2]Department of Toxico/Pharmacology, School of Pharmacy, Shahid Beheshti, University of Medical Sciences, Niayesh Highway, Valiasr Ave, Tehran, Iran

[3]Department of Computational Medicine and Bioinformatics, University of Michigan, Ann Arbor, MI, 48105, USA

[4]Department of Medicine, School of Medicine, McGill University, Montreal, Quebec, Canada

[5]Division of Pulmonary, Critical Care, Sleep and Allergy, Department of Medicine, Miller School of Medicine, University of Miami, Miami, FL, United States

Corresponding author
* Mehdi Mirsaeidi MD, MPH.
Division of Pulmonary, Critical Care, and Sleep
Medicine, University of Miami Miller School of
Medicine, Miami, FL, United States
E-mail: ude.imaim.dem@942msm

### Content 4 suppl figures.

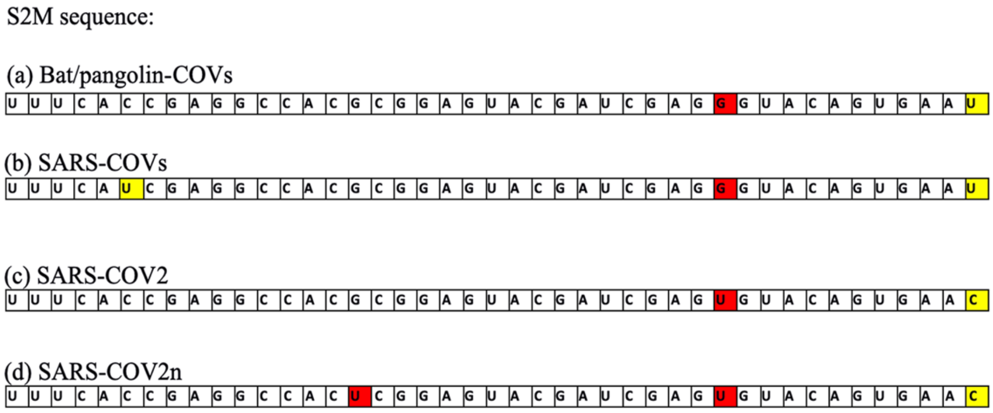


**Figure S1:** A schematic illustration representing the cis-regulatory motifs of the UTR region in SARS-CoV-2 genome (a) bat/pangolin coronaviruses (b) SARS-COVs (c) 32 G-to-U RNA mutations in human SARS-CoV-2 and (d) 16 and 32 G-to-U RNA mutations in human SARS-CoV-2n.

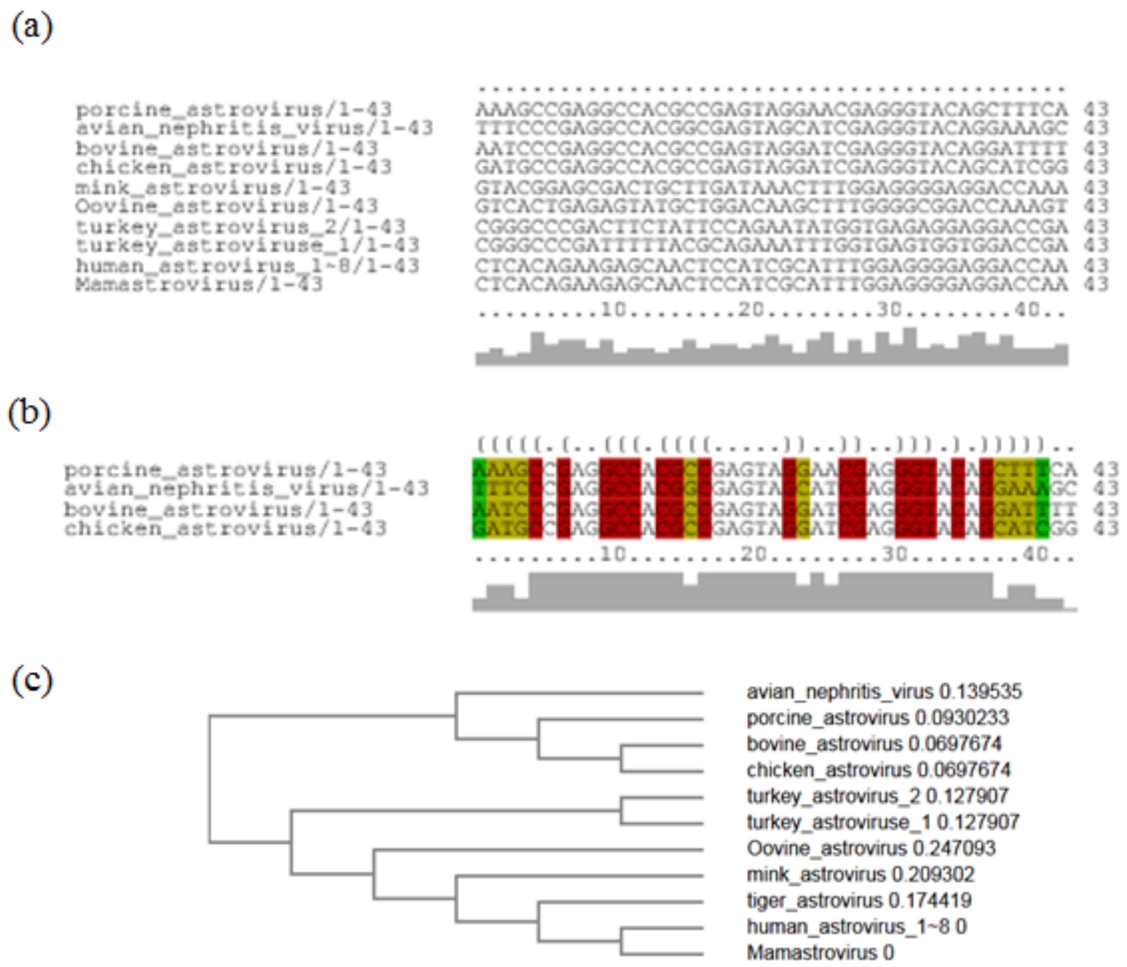


**Figure S2:** Astrovirus S2M sequence motifs. (a) Alignment of stem-forming elements and columns for each genotype. There is one sequence level representation and a second stem loop structure representation. Lines above the alignment indicate stem-forming elements. Columns with Watson Crick pairing changes have been color coded. (b) ClustalW multiple sequence alignment trees display of astroviruses.

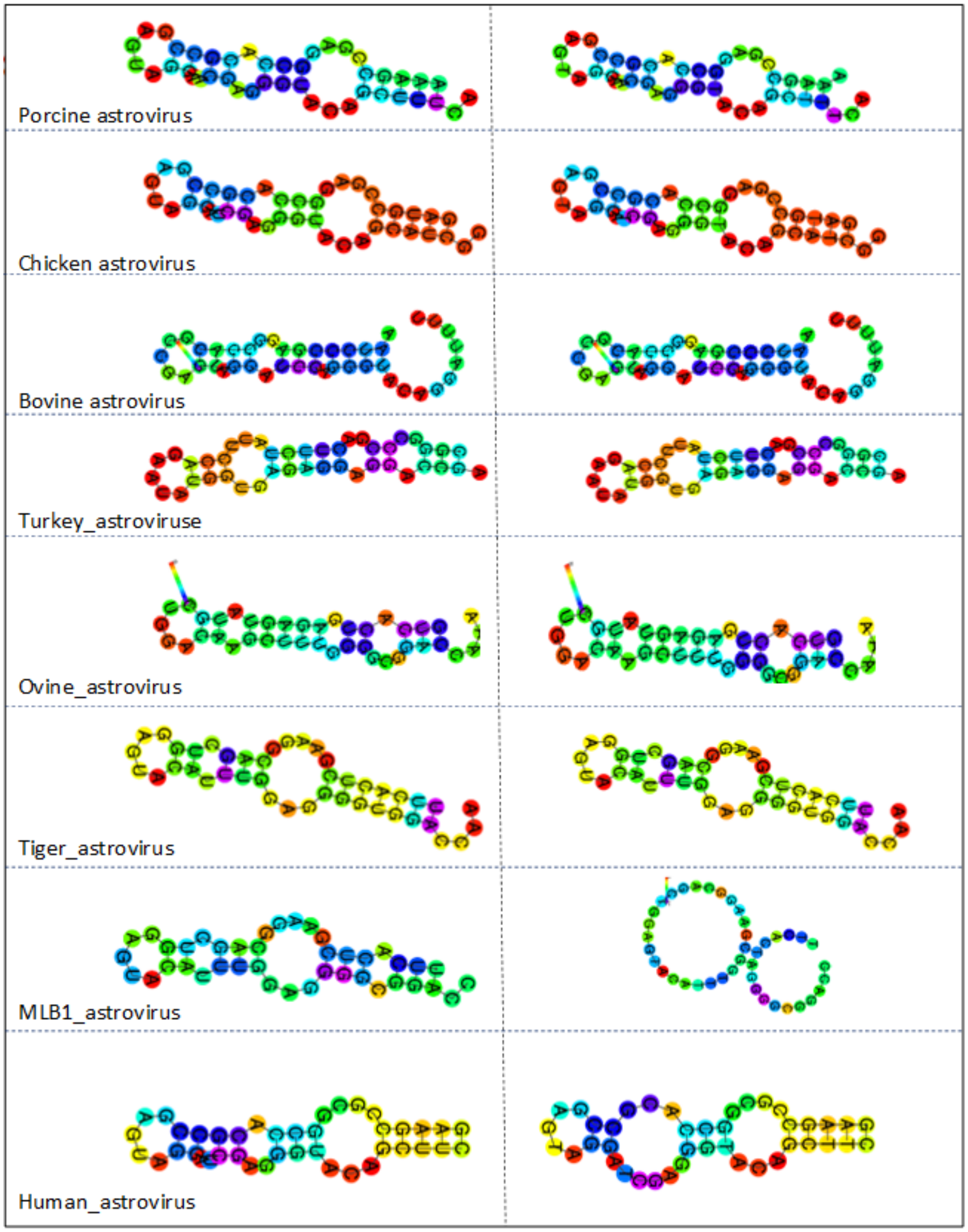


**Figure S3:** S2M sequences were observed in both MFE and centroid secondary structures in coronaviruses. MFE secondary structures are depicted in the left column. Centroid secondary structures are depicted in the right column.

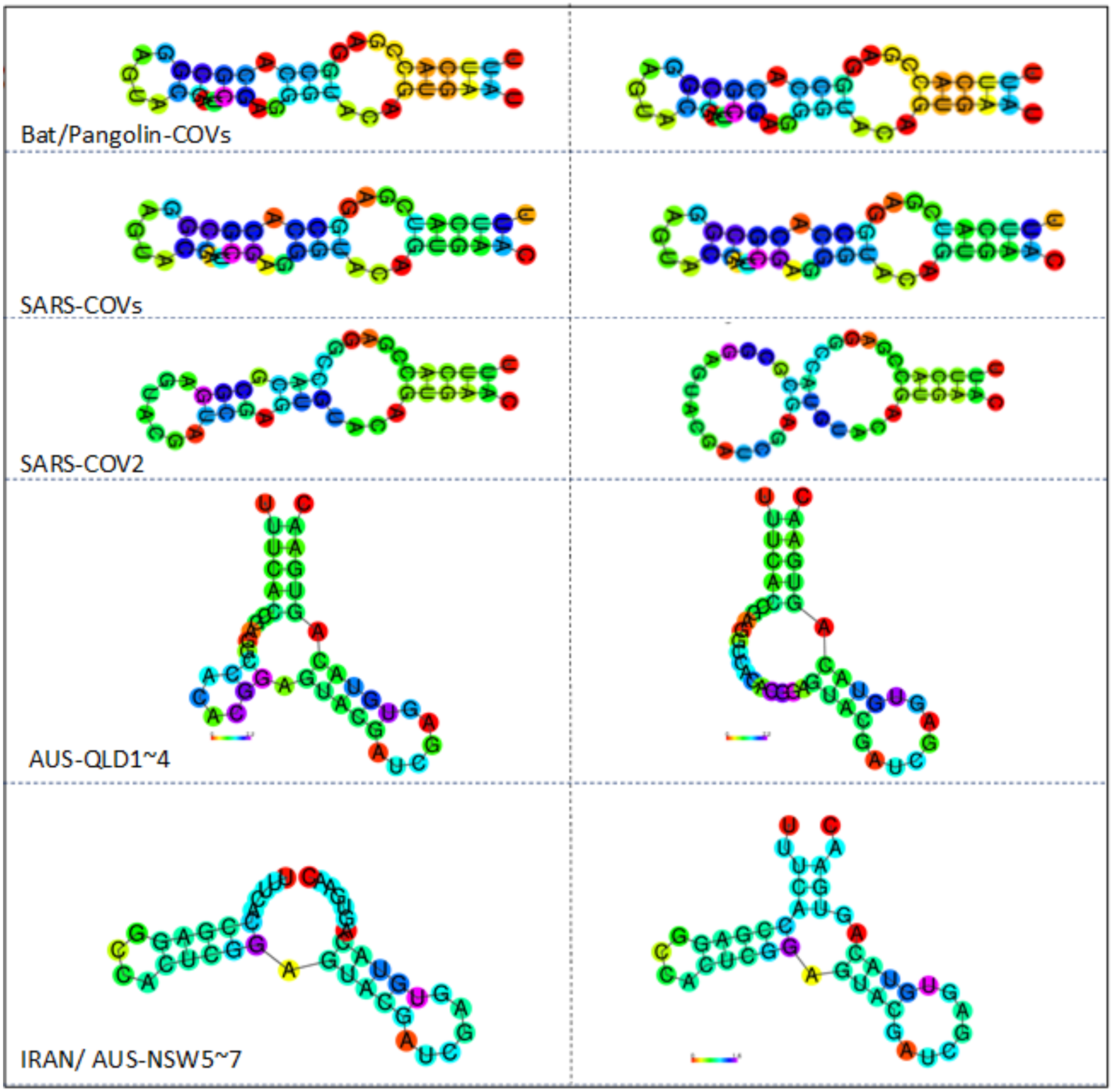


**Figure S4:** S2M sequences were observed in both MFE and centroid secondary structures in astroviruses. MFE secondary structures are depicted in the left column. Centroid secondary structures are depicted in the right column.
